# On the possible inhibition of macrophage inflammatory responses by phytic acid produced by *Echinococcus multilocularis*

**DOI:** 10.1101/2025.09.11.675429

**Authors:** Nicolás Veiga, Anabella A. Barrios, Julia Torres, Álvaro Díaz

## Abstract

*Echinococcus multilocularis*, causing agent of alveolar echinococcosis, down-regulates host immunity ^1,2^. Salzmann *et al* showed that macrophages close to the parasite produce little IL-6 and the parasite extract mimics this inhibition in cultured macrophages ^3^. The authors then showed that *E. multilocularis* contains phytic acid and that commercial phytic acid inhibits cytokine responses in stimulated macrophages. Through comparison with effects of EDTA and intracellular calcium measurements, they concluded that phytic acid inhibits macrophage responses through calcium depletion, extrapolating this conclusion to assays with parasite extract and infection setting. However, the interaction between phytic acid and calcium is complex ^4,5^. Calcium is indeed depleted in the assays with commercial phytic acid according to our quantitative predictions. However additional changes occur including formation of solid calcium phytate, which probably moderates the inhibition of responses observed under calcium depletion. We calculate that in the experiments with parasite extract, phytic acid present cannot deplete calcium and therefore inhibition must be driven by a different mechanism, possibly independent of phytic acid. Further, we argue that calcium depletion is an unlikely *in vivo* immune regulatory mechanism. We propose that biological assays using calcium phytate are needed for ascertaining whether phytic acid has biologically relevant anti-inflammatory activity.

## Main text

Phytic acid refers strictly to the dodeca-acid form of *myo*-inositolhexakisphosphate. Through stepwise deprotonation and/or coordination with metal cations, this compound gives rise to a variety of chemical species. For clarity in nomenclature, phytic acid is represented as H_12_L and the fully deprotonated phytate anion is denoted as L^12-^. In the presence of calcium or magnesium, phytic acid can form both soluble complexes with the general formula [M(H_n_L)]^(10-n)-^ (M being Ca(II) or Mg(II); n = 2-6) and solid salts (commonly referred to as calcium and magnesium phytates) with the formula [M_5_(H_2_L)]·xH_2_O (x = 16 and 22 for Ca and Mg, respectively) ^5,6^. In the present work, we modelled the conditions of the experiments by Salzmann *et al* involving addition of phytic acid and/or EDTA to the macrophage culture media. For this purpose, we used a robust chemical simulation software ^7^ including all the thermodynamic constants for the relevant equilibria involving phytic acid previously determined by us ^5,6^ (more details are available in the Supplementary Materials). The results (Tables 1, S2 and S3) show that in the presence of 1 mM phytic acid, free calcium is indeed depleted. Since additionally the authors showed reduced intracellular calcium levels ^4^, and extracellular calcium is indispensable for macrophage IL-1β and IL-6 responses ^8^, it is clear that calcium depletion is the major factor in the inhibition observed. However, across the experimental conditions used by Salzmann *et al*, the degree of free calcium depletion does not accompany the degree of inhibition of macrophage responses observed. Free calcium depletion by 1 mM phytic acid is essentially complete while 0.25 mM and 0.5 mM EDTA leave 36% and 18% of calcium in free form, respectively (Tables 1, S2 and S3), yet 0.25 mM or 0.5 mM EDTA cause similar or stronger inhibition than 1 mM phytic acid. Along the same lines, calcium depletion is essentially complete using either 1 mM phytic acid alone or 1 mM phytic acid plus 0.5 mM EDTA, but macrophage response inhibition is much more pronounced when EDTA is present. Other two changes predicted to be caused by the addition of 1 mM phytic acid are depletion of free magnesium and acidification of the medium. However, both pH and magnesium depletion are very similar for conditions that inhibit macrophage responses to different degrees, namely 1 mM phytic acid alone and 1 mM phytic acid plus 0.5 mM EDTA (Tables 1, S2 and S3). In addition, macrophage cytokine responses are reported to be enhanced by slight acidosis, and to be inhibited only marginally by low extracellular magnesium^8,9^. The notorious chemical difference between the conditions involving addition of phytic acid alone and the conditions with EDTA (with or without phytic acid) is that only in the former, most (> 85%) of the total calcium is under the form of solid calcium phytate (Tables 1, S2 and S3). The precipitation of calcium phytate may likely have gone undetected due to the small assay volumes used (the expected yield of solid for 0.5 mL is below 0.1 mg). Mammalian cells are known to take up calcium phytate and dephosphorylate phytate in lysosomes ^10^, which makes calcium cation available intracellularly. Therefore, the weaker inhibition of cytokine responses in the absence of EDTA may be rationalized on the basis of solid calcium phytate being taken up by the macrophages and acting as a source of intracellular calcium, thus moderating the effects of extracellular calcium depletion. These deductions are further supported by the observation that, in spite of leaving essentially no free calcium, 1 mM phytic acid inhibits intracellular calcium levels only to the extent of 0.25 mM EDTA ^4^, which as mentioned leaves 36% of total calcium in free form. In sum, whereas extracellular calcium depletion must be a major mechanism underlying inhibition, the observations made are probably influenced by a counteracting effect of calcium phytate formation and its cellular uptake.

**Table 1.**
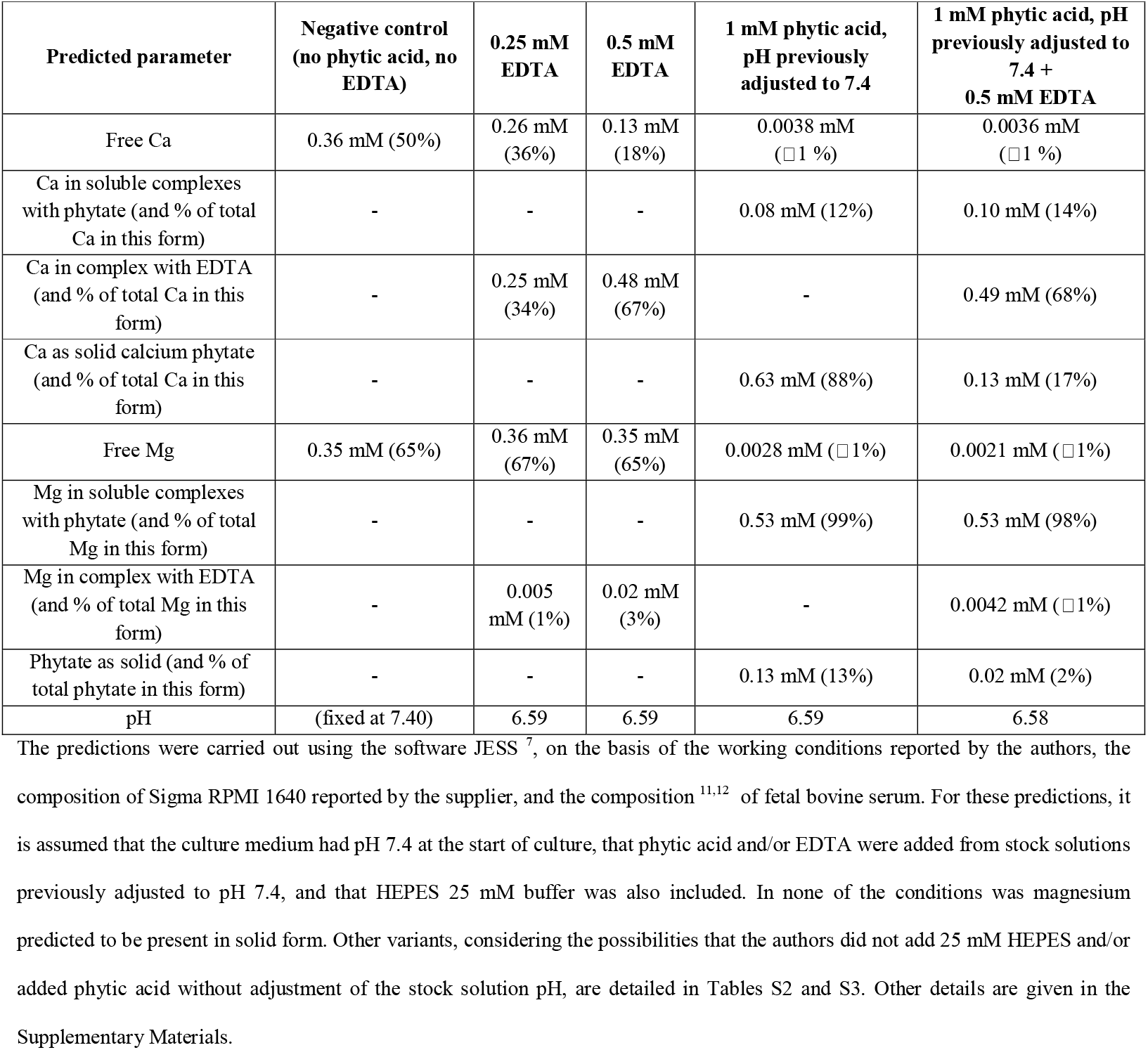
Predictions of the chemical species in which calcium, magnesium and phytic acid are found, as well as pH, in selected *in vitro* experimental conditions used by Salzmann *et al*.

| Predicted parameter | Negative control<br>(no phytic acid, no EDTA) | 0.25 mM EDTA | 0.5 mM EDTA | 1 mM phytic acid, pH previously adjusted to 7.4 | 1 mM phytic acid, pH previously adjusted to 7.4 + 0.5 mM EDTA |
| --- | --- | --- | --- | --- | --- |
| Free Ca | 0.36 mM (50%) | 0.26 mM (36%) | 0.13 mM (18%) | 0.0038 mM (□ 1 %) | 0.0036 mM (□ 1 %) |
| Ca in soluble complexes with phytate (and % of total Ca in this form) | - | - | - | 0.08 mM (12%) | 0.10 mM (14%) |
| Ca in complex with EDTA (and % of total Ca in this form) | - | 0.25 mM (34%) | 0.48 mM (67%) | - | 0.49 mM (68%) |
| Ca as solid calcium phytate (and % of total Ca in this form) | - | - | - | 0.63 mM (88%) | 0.13 mM (17%) |
| Free Mg | 0.35 mM (65%) | 0.36 mM (67%) | 0.35 mM (65%) | 0.0028 mM (□ 1%) | 0.0021 mM (□ 1%) |
| Mg in soluble complexes with phytate (and % of total Mg in this form) | - | - | - | 0.53 mM (99%) | 0.53 mM (98%) |
| Mg in complex with EDTA (and % of total Mg in this form) | - | 0.005 mM (1%) | 0.02 mM (3%) | - | 0.0042 mM (□ 1%) |
| Phytate as solid (and % of total phytate in this form) | - | - | - | 0.13 mM (13%) | 0.02 mM (2%) |
| pH | (fixed at 7.40) | 6.59 | 6.59 | 6.59 | 6.58 |

The predictions were carried out using the software JESS <sup>7</sup>, on the basis of the working conditions reported by the authors, the composition of Sigma RPMI 1640 reported by the supplier, and the composition <sup>11,12</sup> of fetal bovine serum. For these predictions, it is assumed that the culture medium had pH 7.4 at the start of culture, that phytic acid and/or EDTA were added from stock solutions previously adjusted to pH 7.4, and that HEPES 25 mM buffer was also included. In none of the conditions was magnesium predicted to be present in solid form. Other variants, considering the possibilities that the authors did not add 25 mM HEPES and/or added phytic acid without adjustment of the stock solution pH, are detailed in Tables S2 and S3. Other details are given in the Supplementary Materials.

In relation to the experiments involving the parasite extract, we reason that the phytic acid species found in parasite tissues must be saturated with endogenous calcium or magnesium. Which of these cations is phytic acid associated to depends on the cellular compartment where it is present. Any phytic acid present in animal vesicular or extracellular compartments is expected to exist as solid calcium phytate ^6^. This is exemplified by larval *Echinococcus granulosus* (*sensu lato*), which contains extensive deposits of calcium phytate in vesicular compartments and in the acellular laminated layer (LL) ^13–15^. Whereas in the *E. granulosus* LL the calcium phytate deposits are easily detected by electron microscopy, no analogous feature is observed in the *E. multilocularis* LL ^15,16^. Hence the most parsimonious possibility is that the abundant phytic acid discovered by Salzmann *et al* in larval *E. multilocularis* is present as solid calcium phytate in cellular vesicular compartments. On the other hand, any abundant (> 50 μM) phytic acid that may be present in the nuclear or cytosolic compartment is expected to occur as solid magnesium phytate ^5,6^. Upon exposure to a physiological extracellular medium, this compound is expected thermodynamically to give rise to the less soluble solid calcium phytate. Although probably a slow process, this would release magnesium and consume free calcium ^6^. However, even accepting this unlikely scenario, the amounts of phytic acid species present in the experiments with parasite extract are insufficient to significantly deplete calcium. The upper bound for phytic acid content in larval *E. multilocularis* tissue is approximately 0.7% of dry mass, corresponding to the detection limit of a low-sensitivity assay by which phytic acid was detected in larval *E. granulosus* but not in *E. multilocularis* ^13^. Consequently, in the assays using crude parasite extract, the maximum total concentration of phytic acid species present was roughly 0.016 mM (derived from homogenizing 1.16 g of parasite tissue - assumed to contain 10% dry mass - in 7.5 mL, followed by a 1:10 final dilution in cell culture). This amount of phytic acid could remove at most 11% of the total calcium in the system. This figure stems from assuming that all the phytic acid precipitates as [Ca_5_(H_2_L)]·16H_2_O and comparing the calcium thus removed from the solution (0.016 mM x 5 = 0.080 mM) with the total calcium concentration of the system (0.72 mM; see Table S1). It should be noted that the phytic acid concentration in *E. multilocularis* tissues could be considerably lower than the figure used above, as the authors’ data only showed this concentration to be higher than in rodent liver, which contains only 0.0004% phytic acid over dry mass ^4^.

The possibility that phytic acid exerts immune modulatory effects *in vivo* via calcium depletion again encounters the problem that it will be most probably under the form of calcium salt in *E. multilocularis* tissues. In the alternative hypothesis that the parasite stores magnesium phytate in cytosolic/nuclear compartments, the secretion of this compound poses a problem from the perspective of cell biology. If the parasite did manage to secrete magnesium phytate, which as mentioned would slowly consume extracellular calcium, the resulting transient calcium decrease would be followed by a compensatory diffusion of calcium from the surrounding tissues. This would result in a concentration of *total* calcium near the parasite that is higher than that of the surrounding tissues and a concentration of *free* calcium near the parasite that is equal to that found in the extracellular fluids throughout the host’s body. Furthermore, host cells could retrieve calcium by ingesting calcium phytate, as discussed for the *in vitro* experiments. In sum, both in the infection context and in the experiments with parasite extract, free calcium is expected to be present, excluding the possibility to extrapolate to these contexts the observations made under conditions of free calcium depletion (*i*.*e*. experiments with commercial phytic acid).

We conclude that: (i) although calcium depletion is probably the main contributor to the effects of 1 mM phytic acid on macrophages, the observations made are likely influenced by counteracting effects of undetected solid calcium phytate, (ii) the inhibition of macrophage responses observed in the assays with crude parasite extract cannot be explained by calcium depletion and could be unrelated to phytic acid altogether, (iii) calcium depletion by phytic acid is not a plausible immune regulation mechanism *in vivo*. The possibility that calcium phytate particles inhibit macrophage responses under conditions of physiological extracellular concentrations of free calcium certainly cannot be ruled out. The inflammatory response to particles of the *E. granulosus* LL injected intraperitoneally is moderated by its calcium phytate component, through unknown mechanisms ^17^. Calcium phytate, not commercially available, can be synthesized from commercial phytic and a soluble calcium salt ^6^. *In vitro* experiments in which macrophages are confronted with this solid compound entail no risk of metal ion depletion or acidification, and should be informative.

## Author contributions

Nicolás Veiga: Investigation - chemical data modeling, Writing – review & editing; Anabella A. Barrios: Investigation - biological data curation, Writing – review & editing; Julia Torres: Conceptualization, Writing – review & editing; Álvaro Díaz: Conceptualization, Writing – original draft, review & editing.

## Data availability statement

Data shown in Tables 1, S2 and S3 were obtained by a process described in full detail in the Supplementary section.

## References

1. Brunetti, E. & White, A. C. Cestode infestations: hydatid disease and cysticercosis. Infect Dis Clin North Am 26, 421–435 (2012).

2. Wang, L. et al. TIM4+macrophages suppress the proinflammatory response to maintain the chronic alveolar echinococcosis infection. Front Cell Infect Microbiol 15, 1600686 (2025).

3. Ou, Z. et al. Spatiotemporal Transcriptomic Profiling Reveals the Dynamic Immunological Landscape of Alveolar Echinococcosis. Adv Sci (Weinh) 12, e2405914 (2025).

4. Salzmann, M. et al. Phytic acid impairs macrophage inflammatory response in Echinococcus multilocularis infection. Commun Biol 8, 871 (2025).

5. Torres, J. et al. Solution behaviour of myo-inositol hexakisphosphate in the presence of multivalent cations. Prediction of a neutral pentamagnesium species under cytosolic/nuclear conditions. Journal of Inorganic Biochemistry 99, 828–840 (2005).

6. Veiga, N. et al. The behaviour of myo-inositol hexakisphosphate in the presence of magnesium(II) and calcium(II): Protein-free soluble InsP6 is limited to 49μM under cytosolic/nuclear conditions. Journal of Inorganic Biochemistry 100, 1800–1810 (2006).

7. May, P. M. & Filella, M. Open Access to the JESS Chemical Reaction Database. J Solution Chem 52, 1149–1152 (2023).

8. Libako, P., Nowacki, W., Castiglioni, S., Mazur, A. & Maier, J. A. M. Extracellular magnesium and calcium blockers modulate macrophage activity. Magnes Res 29, 11–21 (2016).

9. Erra Díaz, F., Dantas, E. & Geffner, J. Unravelling the Interplay between Extracellular Acidosis and Immune Cells. Mediators Inflamm 2018, 1218297 (2018).

10. Windhorst, S. et al. Tumour cells can employ extracellular Ins(1,2,3,4,5,6)P(6) and multiple inositol-polyphosphate phosphatase 1 (MINPP1) dephosphorylation to improve their proliferation. Biochem J 450, 115–125 (2013).

11. Price, P. J. & Gregory, E. A. Relationship between in vitro growth promotion and biophysical and biochemical properties of the serum supplement. In Vitro 18, 576–584 (1982).

12. An, L. et al. Magnesium is a critical element for competent development of bovine embryos. Theriogenology 140, (2019).

13. Irigoín, F., Ferreira, F., Fernández, C., Sim, R. B. & Díaz, A. myo-Inositol hexakisphosphate is a major component of an extracellular structure in the parasitic cestode Echinococcus granulosus. 8 (2002).

14. Casaravilla, C. et al. Characterization of myo-inositol hexakisphosphate deposits from larval Echinococcus granulosus. FEBS Journal 273, 3192–3203 (2006).

15. Irigoín, F. et al. Unique precipitation and exocytosis of a calcium salt of myo - inositol hexakisphosphate in larval Echinococcus granulosus. J. Cell. Biochem. 93, 1272–1281 (2004).

16. Ingold, K., Gottstein, B. & Hemphill, A. High molecular mass glycans are major structural elements associated with the laminated layer of in vitro cultivated Echinococcus multilocularis metacestodes. Int J Parasitol 30, 207–214 (2000).

17. Grezzi, L., González, C., Díaz, Á. & Casaravilla, C. The Acute Inflammatory Potential of Particles From the Echinococcus granulosus Laminated Layer Is Moderated by Its Calcium Inositol Hexakisphosphate Component. Parasite Immunol 46, e13040 (2024).

