## Supplemental Materials for "On the possible inhibition of macrophage inflammatory responses by phytic acid produced by *Echinococcus multilocularis*"

**Supplementary Materials**

Simulation of the chemical speciation

The distribution of chemical species was determined using the Joint Expert Speciation System (JESS), a tool for modelling chemical speciation in complex environments ^1^. JESS incorporates an extensive thermodynamic database (>280,000 equilibrium constants) covering reactions of biological, environmental, and technological relevance. The equilibrium constants for the protonation, complexation and precipitation of phytic acid species were added to the JESS database from our previous works ^2,3^.

For each calculation, JESS selects a thermodynamically consistent set of equations and solves them iteratively to obtain the chemical species distribution. All simulations were performed at 37 °C, 1 atm, and with an oxygen redox potential equivalent to 18.2% O_2_ (fixed p*E* = 13.2), similar to that prevalent in cell culture incubators ^4^. Ionic strength was calculated at equilibrium. The precipitation of solid phases was tested iteratively, from higher to lower supersaturation indices. The precipitation of apatites (Ca_5_X(PO^­^_4_)_3_(s) with X = Cl^−^, F^−^, OH^−^) was omitted due to kinetic constraints ^5^.

We followed a three-step procedure:

*Step 1: Reference simulations*

We first modeled the two control media: RPMI 1640 with 10% v/v FBS, either without or with 25 mM HEPES (sodium salt). Table S1 lists the chemical components and the corresponding millimolar concentrations considered in these simulations. These components were selected based on the reported composition of Sigma RPMI 1640 medium, the general composition of fetal bovine serum (FBS) ^6,7^ and the possible supplementation with HEPES. Four quantitatively minor low molecular mass components (creatinine, bilirubin, choline and *myo*-inositol), as well as protein components, were excluded due to insufficient thermodynamic data to reliably model their distribution. The final thermodynamic database used for the simulations contained 1,133 equilibria and 10,578 equilibrium constants.

For the two reference simulations, pH was fixed at 7.4. JESS then calculated the total concentration of protons, which we kept as reference values ([H^+^]_ctrl_ and [H^+^]_ctrl+HEPES_) for later simulations. Notably, the resulting CO_2_(g) partial pressure in equilibrium with both control simulations was 0.04 atm (4%), in close agreement with the value at which most cell culture incubators are set (0.05 atm; 5%). This agreement supports the reliability of the model.

**Table S1. Chemical components considered in the simulations and their milimolar concentrations included for the control conditions.**

| Component | RPMI+FBS (pH = 7.4) | RPMI+FBS + HEPES  (pH = 7.4) |
| --- | --- | --- |
| Na^+^ | 137.87 | 162.88 |
| K^+^ | 5.95 | 5.95 |
| Mg^2+^ | 0.538 | 0.538 |
| Ca^2+^ | 0.720 | 0.720 |
| Cl^−^ | 108.12 | 108.12 |
| PO_4_^3−^ | 5.39 | 5.39 |
| CO_3_^2-^ | 21.43 | 21.43 |
| urea | 0.571 | 0.571 |
| urate | 0.0173 | 0.0173 |
| glucose | 10.68 | 10.68 |
| L-Arginine | 1.033 | 1.033 |
| L-Asparagine | 0.340 | 0.340 |
| L-Aspartic | 0.135 | 0.135 |
| L-Cystine | 0.187 | 0.187 |
| L-Glutamic | 0.122 | 0.122 |
| L-Glutamine | 1.848 | 1.848 |
| Glycine | 0.120 | 0.120 |
| L-Histidine | 0.087 | 0.087 |
| Hydroxy-L-Proline | 0.137 | 0.137 |
| L-Isoleucine | 0.343 | 0.343 |
| L-Leucine | 0.343 | 0.343 |
| L-Lysine | 0.197 | 0.197 |
| L-Methionine | 0.0904 | 0.0904 |
| L-Phenylalanine | 0.0817 | 0.0817 |
| L-Proline | 0.156 | 0.156 |
| L-Serine | 0.257 | 0.257 |
| L-Threonine | 0.151 | 0.151 |
| L-Tryptophan | 0.0220 | 0.0220 |
| L-Tyrosine | 0.0993 | 0.0993 |
| L-Valine | 0.154 | 0.154 |
| D-Biotin | 0.000737 | 0.000737 |
| Folate | 0.00204 | 0.00204 |
| Niacinamide | 0.00737 | 0.00737 |
| p-Amino benzoate | 0.00656 | 0.00656 |
| Glutathione | 0.00293 | 0.00293 |
| HEPES (anionic form) | 0 | 25 |

*Step 2: Simulations for calculating proton contributions from added phytic acid or EDTA pre-adjusted to pH 7.4*

Next, we calculated the proton concentrations contributed by the addition of 1 mM phytic acid or Na₂H₂EDTA at 0.25 o 0.5 mM (in all cases adjusted to pH 7.4 before addition). These gave the values [H^+^]_phytic_, [H^+^]_EDTA0.25_ and [H^+^]_EDTA0.5_ respectively, which were retained for later use.

*Step 3: Final simulations*

We simulated the possibilities of addition of EDTA 0.25 mM, EDTA 0.5 mM, and phytic acid 1 mM plus EDTA 0.5 mM. In addition to the variants with and without HEPES, we considered the possibilities that phytic acid was added without prior pH adjustment (as dodecaprotonated acid, H_12_L or as pentasodium salt, Na_5_H_7_L), or was added after adjustment to pH 7.4 (in which case it does not matter whether it was initially H_12_L or Na_5_H_7_L). EDTA was considered in all cases to have been added from a stock solution with prior pH adjustment to 7.4.

So, for setting up the final simulations, all components of each control medium were included at the corresponding reference concentration values (Table S1). In addition, phytate (L^12-^) and EDTA were included as components, at the corresponding concentration depending on the specific simulation. Finally, for the H^+^ component, the initial total proton concentration was set as [H^+^]_ctrl_ or [H^+^]_ctrl+HEPES_ (step 1), then adding the proton contributions from the added phytic acid or EDTA. In detail, on one hand, for simulations with the addition of solutions with prior pH adjustment to pH 7.4, [H^+^]_phytic_ and/or [H^+^]_EDTA0.25_/[H^+^]_EDTA0.5_ (step 2) were added. On the other hand, for simulations in which phytic acid was added without prior pH adjustment (as H_12_L or Na_5_H_7_L), the corresponding proton loads (12 or 7 mM, respectively) were included directly. In all cases, pH was calculated as an output of the simulation.

Supplementary predictions

The following tables show the results on the principal chemical species in which calcium, magnesium and phytic acid are found, as well as pH, in selected *in vitro* experimental conditions explored. These data complement those in Table 1 by taking into account experimental variants that were not detailed in the article by Salzmann *et al*.

**Table S2. Results of chemical speciation assuming that HEPES buffer was not used.**

| **Predicted parameter** | **Negative control (no phytic acid, no EDTA)** | **0.25 mM EDTA** | **0.5 mM EDTA** | **1 mM phytic acid, pH previously adjusted to 7.4** | **1 mM phytic acid, pH previously adjusted to 7.4 +**  **0.5 mM EDTA** |
| --- | --- | --- | --- | --- | --- |
| Free Ca (and % of total Ca in this form) | 0.34 mM (48%) | 0.26 mM (36%) | 0.13 mM (18%) | 0.0040 mM (˂1%) | 0.0039 mM (˂1%) |
| Ca in soluble complexes with phytate (and % of total Ca in this form) | - | - | - | 0.08 mM (12%) | 0.11 mM (15%) |
| Ca in complex with EDTA (and % of total Ca in this form) | - | 0.25 mM (34%) | 0.48 mM (67%) | - | 0.48 mM (67%) |
| Ca as solid (and % of total Ca in this form) | - | - | - | 0.63 mM (87%) | 0.12 mM (17%) |
| Free Mg | 0.34 mM (63%) | 0.36 mM (66%) | 0.35 mM (65%) | 0.0034 mM (˂1%) | 0.0026 mM (˂1%) |
| Mg in soluble complexes with phytate (and % of total Mg in this form) | - | - | - | 0.53 mM (99%) | 0.53 mM (99%) |
| Mg in complex with EDTA (and % of total Mg in this form) | - | 0.005 mM (1%) | 0.02 mM (3%) | - | 0.0048 mM (1%) |
| Phytate as solid (and % of total phytate in this form) | - | - | - | 0.13 mM (13%) | 0.02 mM (2%) |
| pH | (fixed at 7.40) | 6.50 | 6.50 | 6.49 | 6.49 |

These predictions were carried out analogously to those shown in Table 1, except that buffer HEPES 25 mM was not included. In none of the conditions was magnesium predicted to be present in solid form.

**Table S3. Results of chemical speciation considering two possibilities for the addition of commercial phytic acid without prior pH adjustment.**

| **Addition of HEPES** | **Predicted parameter** | **1 mM phytic acid** | **1 mM phytic acid (penta-sodium salt)** | **1 mM phytic acid +**  **0.5 mM EDTA** | **1 mM phytic acid (penta-sodium salt) +**  **0.5 mM EDTA** |
| --- | --- | --- | --- | --- | --- |
| **No** | Free Ca (and % of total Ca in this form) | 0.005 mM (˂1%) | 0.0042 mM (˂1%) | 0.0048 mM (˂1%) | 0.0041 mM (˂1%) |
|  | Ca in soluble complexes with phytate (and % of total Ca in this form) | 0.09 mM (13%) | 0.08 mM (12%) | 0.11 mM (15%) | 0.11 mM (15%) |
|  | Ca in complex with EDTA (and % of total Ca in this form) | - | - | 0.48 mM (67%) | 0.48 mM (67%) |
|  | Ca as solid (and % of total Ca in this form) | 0.62 mM (86%) | 0.63 mM (87%) | 0.12 mM (16%) | 0.12 mM (17%) |
|  | Free Mg | 0.005 mM (˂1%) | 0.0037 mM (˂1%) | 0.0038 mM (˂1%) | 0.0028 mM (˂1%) |
|  | Mg in soluble complexes with phytate (and % of total Mg in this form) | 0.53 mM (99%) | 0.53 mM (99%) | 0.52 mM  (98%) | 0.53 mM (98%) |
|  | Mg in complex with EDTA (and % of total Mg in this form) | - | - | 0.0056 mM (1%) | 0.0049 mM (1%) |
|  | Phytate as solid (and % of total phytate in this form) | 0.12 mM (12%) | 0.13 mM (13%) | 0.02 mM (2%) | 0.02 mM (2%) |
|  | pH | 6.35 | 6.46 | 6.34 | 6.46 |
| **Yes** | Free Ca | 0.004 mM (˂1%) | 0.0039 mM (˂1%) | 0.0039 mM (˂1%) | 0.0038 mM (˂1 %) |
|  | Ca in soluble complexes with phytate (and % of total Ca in this form) | 0.08 mM (11%) | 0.08 mM (12%) | 0.10 mM (14%) | 0.10 mM (14%) |
|  | Ca in complex with EDTA (and % of total Ca in this form) | - | - | 0.49 mM (67%) | 0.49 mM (68%) |
|  | Ca as solid (and % of total Ca in this form) | 0.63 mM (87%) | 0.63 mM (88%) | 0.12 mM (17%) | 0.12 mM (17%) |
|  | Free Mg | 0.0031 (˂1%) | 0.0029 mM (˂1%) | 0.0024 mM (˂1%) | 0.0022 mM (˂1%) |
|  | Mg in soluble complexes with phytate (and % of total Mg in this form) | 0.53 mM (99%) | 0.53 mM (99%) | 0.53 mM (98%) | 0.53 mM (98%) |
|  | Mg in complex with EDTA (and % of total Mg in this form) | - | - | 0.0044 mM (˂1%) | 0.0042 mM (˂1%) |
|  | Phytate as solid (and % of total phytate in this form) | 0.13 mM (13%) | 0.13 mM (13%) | 0.02 mM (2%) | 0.02 mM (2%) |
|  | pH | 6.54 | 6.57 | 6.53 | 6.57 |

These predictions were carried out analogously to those shown in Table 1, except that for each of these conditions, variants cover the possibilities that the authors added Sigma product 593648 (*bona fide* phytic acid solution) or Sigma product P8810 (sodium salt, which according to the supplier analysis is predominantly a penta-sodium salt) without adjustment of the stock solution pH. Variants are also given to cover the possibilities that the authors added or did not add 25 mM HEPES to the culture medium. In none of the conditions was magnesium predicted to be present in solid form.

**Supplementary Material References**

1.	May, P. M. & Filella, M. Open Access to the JESS Chemical Reaction Database. *J Solution Chem* **52**, 1149–1152 (2023).

2.	Veiga, N. *et al.* The behaviour of myo-inositol hexakisphosphate in the presence of magnesium(II) and calcium(II): Protein-free soluble InsP6 is limited to 49μM under cytosolic/nuclear conditions. *Journal of Inorganic Biochemistry* **100**, 1800–1810 (2006).

3.	Torres, J. *et al.* Solution behaviour of myo-inositol hexakisphosphate in the presence of multivalent cations. Prediction of a neutral pentamagnesium species under cytosolic/nuclear conditions. *Journal of Inorganic Biochemistry* **99**, 828–840 (2005).

4.	Wenger, R. H., Kurtcuoglu, V., Scholz, C. C., Marti, H. H. & Hoogewijs, D. Frequently asked questions in hypoxia research. *Hypoxia (Auckl)* **3**, 35–43 (2015).

5.	Boskey, A. L. & Posner, A. S. Conversion of amorphous calcium phosphate to microcrystalline hydroxyapatite. A pH-dependent, solution-mediated, solid-solid conversion. *J. Phys. Chem.* **77**, 2313–2317 (1973).

6.	Price, P. J. & Gregory, E. A. Relationship between in vitro growth promotion and biophysical and biochemical properties of the serum supplement. *In Vitro* **18**, 576–584 (1982).

7.	An, L. *et al.* Magnesium is a critical element for competent development of bovine embryos. *Theriogenology* **140**, (2019).
